# Cross-species brain-wide mapping reveals a conserved and coordinated network engaged by NAc DBS

**DOI:** 10.1101/2024.09.08.611940

**Authors:** Austin Y. Feng, Daniel A. N. Barbosa, Austen B. Casey, Daniel R. Rijsketic, Juliana S. Salgado, Harvey Huang, Robert C. Malenka, Dora Hermes, Kai J. Miller, Casey H. Halpern, Boris D. Heifets

**Author notes:** These authors contributed equally: Austin Y. Feng, Daniel A. N. Barbosa. These senior authors contributed equally: Casey H. Halpern, Boris D. Heifets. Lead contact: Boris D. Heifets.

## Abstract

**Summary:** Nucleus accumbens (NAc) deep brain stimulation (DBS) has been increasingly explored as a treatment modality for refractory neuropsychiatric disorders. Uncovering the accumbens network that is engaged by DBS is a critical step forward in understanding how modulating this important node impacts the broader mesocorticolimbic circuit. Using whole-brain clearing and unbiased, brain-wide neural activity mapping, we found that NAc DBS increases neural activity in a coordinated mesocorticolimbic network in mice. Simultaneous intracranial electrophysiology recordings from the human NAc and brief stimulation epochs of homologous mesocorticolimbic nodes revealed similar connectivity. Altogether, these results identify specific connectivity conserved across species within the mesocorticolimbic circuit that may underlie mechanisms of NAc DBS.

## Main

Maladaptive reward-related behaviors are pervasive in multiple public health crises, including depression, substance use disorders, and eating disorders, which together have a global lifetime prevalence of nearly 40 percent.^1^ Over 10% of such patients have intractable disorders, leading to tremendous economic and quality of life burdens, underscoring the need for novel therapeutic approaches.^2^ Despite the broad range of pathophysiology represented in such appetitive and affective disorders, they are linked by profound deficits in reward processing that may be amenable to modulation via novel therapeutic approaches transdiagnostically.^3,4^ The nucleus accumbens (NAc), a major hub for reward processing in the brain, is located within the ventral striatum where it makes key connections to the orbito-frontal and medial prefrontal cortices, basolateral amygdala, hippocampus, ventral tegmental area and ventral pallidum.^5–8^ These circuits have been heavily implicated in the motivational guidance of actions, hedonic value association, integration of contextual information, executive control, and reward acquisition.^9–15^ Both clinical and preclinical data implicate dysregulated NAc circuitry in anhedonia, eating disorders, and compulsive reward seeking in substance use disorders, and eating disorders.^3,7,16–18^ Given these important functions, the NAc increasingly draws attention as a promising neuromodulation target for refractory neuropsychiatric conditions.

Deep brain stimulation (DBS) has been effectively utilized to modulate neural circuits, most notably for the treatment of movement disorders^19–21^, and a growing body of evidence support the use of DBS to modulate circuits underlying neuropsychiatric illnesses and related conditions. In particular, DBS of the NAc region has demonstrated promise for a wide range of indications, including substance use disorder, obsessive-compulsive disorder, major depressive disorder, binge-eating disorder, and obesity.^22–29^ NAc DBS has been found to suppress drug seeking and loss of control eating behaviors as well as induce antidepressant and anxiolytic responses in rodent models of addiction, binge-eating/drinking, and anxiety/depression.^15,16,30–32^ These findings highlight the potential efficacy of NAc DBS, however, major gaps remain in our understanding of the network dynamics underlying this complex intervention. Clinical outcomes following DBS of the NAc region are notably variable, highlighting an inadequate understanding of its therapeutical mechanisms.^33,34^ A fresh look at neural network engagement associated with NAc DBS would benefit from newly available methods for broad-based, unbiased analysis of network activity. Previous attempts at mapping long-distance neural effects of NAc DBS in mice have been limited by several factors, including constrained anatomical planes of investigation, restricted volumes for analysis, as well as cell counting reliability across sections in traditional 2D immunohistochemistry techniques.^30^ Moreover, methodological discrepancies for appropriately bridging human and non-human stimulation data have substantially limited the translatability of prior investigations.^35^ Here, we mapped the brain-wide effects of NAc DBS on neural activity in mice, defined by expression of the immediate early gene c-Fos, using immunolabeling-enabled 3D imaging of solvent-cleared organs (iDISCO) which accurately preserves neuroanatomical structures of the brain across macro and microscopic scales.^36^ In conjunction with an unbiased, supervised machine-learning-based method for pixel classification and cell counting, for the first time, we performed an immunohistochemical examination of whole brains undergoing NAc DBS in a previously unattainable manner.^37^ We then tested the translatability of key circuit-hypotheses based on these findings in a human subject with rare direct recordings of evoked potentials in the NAc in response to electrical stimulation of multiple mesocorticolimbic regions implanted with depth electrodes.

## Results

### Increased activity of discrete mesocorticolimibic neuron clusters with NAc DBS

We analyzed brain-wide neural activity in mice, indexed by Fos expression, in response to bilateral NAc DBS. 8-week-old male and female mice were implanted bilaterally with bipolar electrodes in the NAc medial shell using validated methods (Fig. 1A).^16,38^ All mice received either active or sham DBS delivered as continuous biphasic pulses (130 Hz, 0.1 mA, 90 µs) in their home cage environment, followed by brain extraction and fixation (Fig. 1B). Stimulation parameters were chosen for their efficacy at modulating maladaptive mouse behaviors.^15,16,38^ Fixed brains were immunolabeled, cleared via iDISCO, and imaged with light sheet fluorescence microscopy (LSFM; [Fig. 1C-F]). This iDISCO procedure preserves the spatial relationship between regions compared to the reconstruction of brain slices with conventional immunohistochemistry (Fig. 1C).^36^ Image preprocessing, automation, and visualization was facilitated with an initial version of UN-biased high-Resolution Analysis and Validation of Ensembles using Light sheet images (UNRAVEL; http://github.com/b-heifets/UNRAVEL) and custom python scripts.^39^ Uniformity of image intensities was achieved through background subtraction with a rolling-ball algorithm (Fig. 1D). We registered samples to a standard atlas using the multi-modal image registration and connectivity analysis (MIRACL) pipeline, substituting an averaged template brain generated with iDISCO+ and LSFM in place of one obtained via serial two-photon microscopy, to better match our samples’ autofluorescence profile (Fig. 1E). ^40,41^ Immunofluorescence images were pre-processed and warped to atlas space and spatially smoothed (50 µm kernel) for voxel-wise analyses. Using a generalized linear model (GLM) of voxel intensity, nonparametric permutation inference testing was performed using the “*randomise”* tool in the FMRIB Software Library (FSL) to identify areas of differential Fos expression between active and sham cohorts (Fig. 1F). In addition to the NAc region that received direct stimulation, increased voxel intensities were observed across a distributed network of cortical and subcortical structures for the active DBS cohort (Fig. 1G), including areas within the anterior olfactory nucleus (AON), lateral orbital area (ORBI), gustatory area (GUST), anterior cingulate area (ACA), basolateral amygdala (BLA), ventral hippocampus (vHP), and periaqueductal gray (PAG), as well as scattered portions of the secondary motor area (MOs), primary somatosensory areas (SSp), rostral lateral septal nucleus (LSr), and agranular insular area (AI). Decreased voxel intensities were observed in very small portions of the primary motor area (MOp) and superior colliculus (SC).

**Figure 1.**
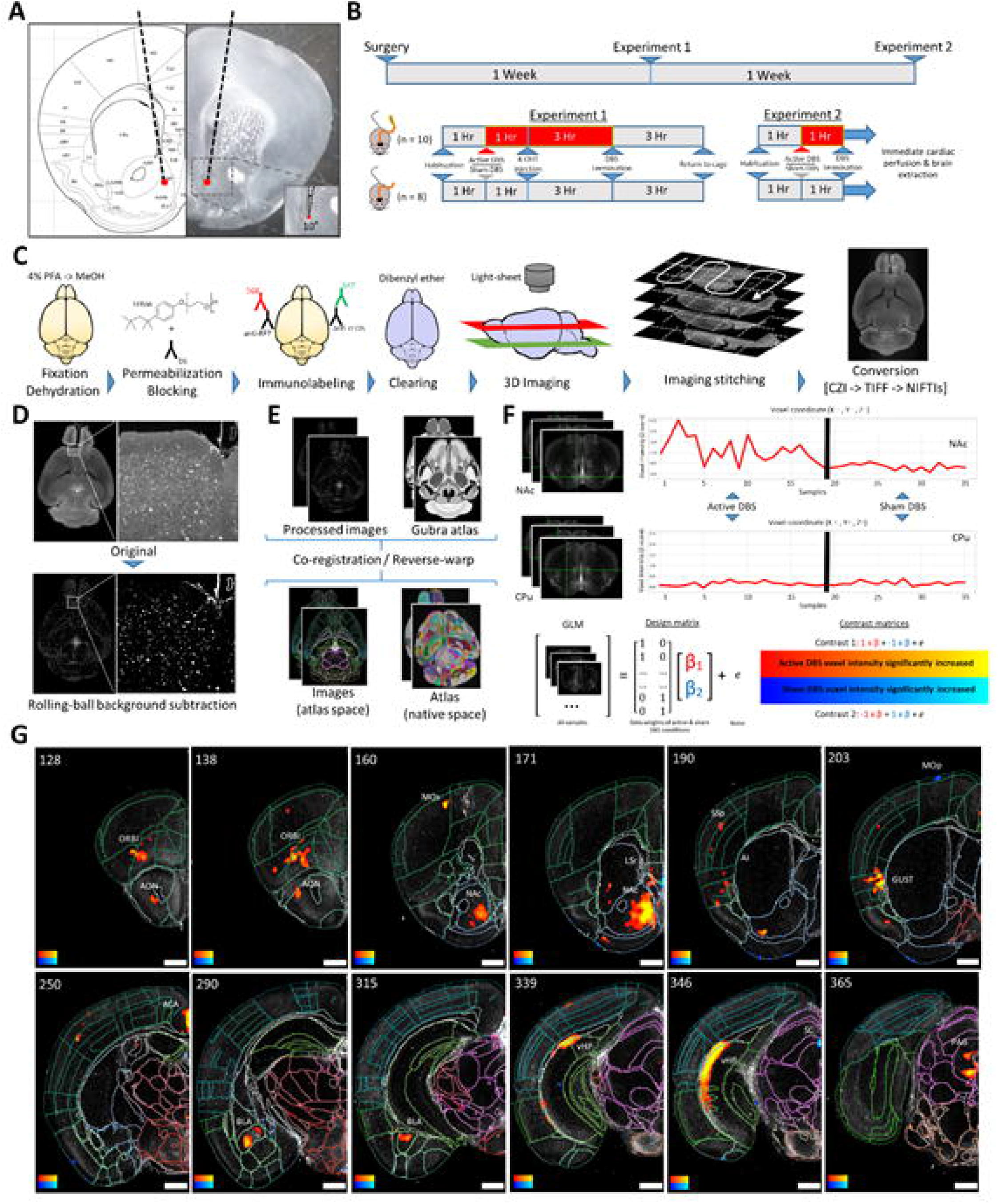
Unbiased, brain-wide assessment of changes in Fos expression induced by NAc DBS. **A**. Nucleus accumbens (NAc) target coordinates – AP: +1.34 mm from bregma, LM: -/+ 0.6 mm, DV: +4.25 mm (within shell). Electrodes were angled 10° relative to the target to facilitate bilateral implantation. **B**. Schematic for NAc DBS experiments. Mice were randomly assigned to either active (n = 10) or sham (n = 8) DBS cohorts. Each cohort had an equal number of male and female mice. **C**. Summary of iDISCO and LSFM pipeline. See *Methods* for details. **D**. Preprocessing with rolling-ball background subtraction (radius = 4 pixels) to remove autofluorescence and normalize voxel intensities within and across samples **E**. Analysis pipeline overview. Co-registration was first performed to generate the transformation matrices of the mouse brain atlas from warping the average template to each sample. Next, inverse deformation fields from this were used to warp immunofluorescence images into atlas space. After smoothing, voxel-wise permutation testing was performed. **F**. Assessing voxel-wise differences in Fos expression. GLM represents voxel intensities of concatenated sequences of active and sham samples as a linear combination of predictors (beta weights β1 and β2 for active and sham DBS, respectively) and noise. Nonparametric permutation thresholding is accomplished with the FSL “*randomise”* tool to define voxels with different Fos expression between groups. Voxels are resampled and a distribution of permuted t-statistics are generated. Rejection of null hypothesis (complete observation exchangeability) depends on the t statistic’s percentile in the permutation distribution. For instance, greater intensities are observed in the example voxel within the NAc in the active versus sham DBS cohorts. In comparison, similar voxel intensities are observed in the voxel within the CPu across both cohorts. **G**. Initial nonparametric permutation inference testing findings. Active NAc DBS stimulates multiple non-contiguous neuronal populations. Brain slices with LSFM atlas overlays (edges) are presented coronally. Warm and cool color scales, respectively, indicate significantly increased or decreased voxel intensities for active and sham DBS cohorts; both spectrums range between t statistics of 2.75 and 4.0, increasing from left to right on the intensity legend. Coronal slice numbers are indicated. NAc, nucleus accumbens; DBS, deep brain stimulation; AP, anterior-posterior; LM, lateral-medial; DV, dorsal-ventral, iDISCO, immunolabeling-enabled three-dimensional imaging of solvent-cleared organs; GLM, generalized linear model; CPu, dorsal striatum; LSFM, light sheet fluorescence microscopy; AON, anterior olfactory nucleus; ORBl, lateral orbital area; NAc, nucleus accumbens; GUST, gustatory area; ACA, anterior cingulate area; BLA, basolateral amygdala; vHP, ventral hippocampus; PAG, periaqueductal gray; MOp, primary motor area; MOs, secondary motor area; SSp, primary somatosensory area; LSr, rostral lateral septal nucleus; AI, agranular insular area; SC, superior colliculus.

Based on the between-group differences in voxel intensity, we then identified four discrete clusters with increased activity that were non-contiguous to the site of stimulation. This modulation occurred in the BLA (Fig. 2A, left), vHP (Fig. 2B, left), lateral orbitofrontal cortex (LOFC; Fig. 2C, left), and GUST (Fig. 2D, left). As predicted, another cluster of increased voxel intensity expression was identified in the NAc, specifically in the area surrounding the electrode tips (Fig. 2E, left).^16^ These five clusters represented areas of greater voxel intensity in the active versus sham cohorts, suggesting increased Fos expression in the active NAc DBS cohort (Fig. 2F). No discrete clusters of voxels with decreased intensity were identified for further analyses (see *Online Methods* for details on clustering approach).

**Figure 2.**
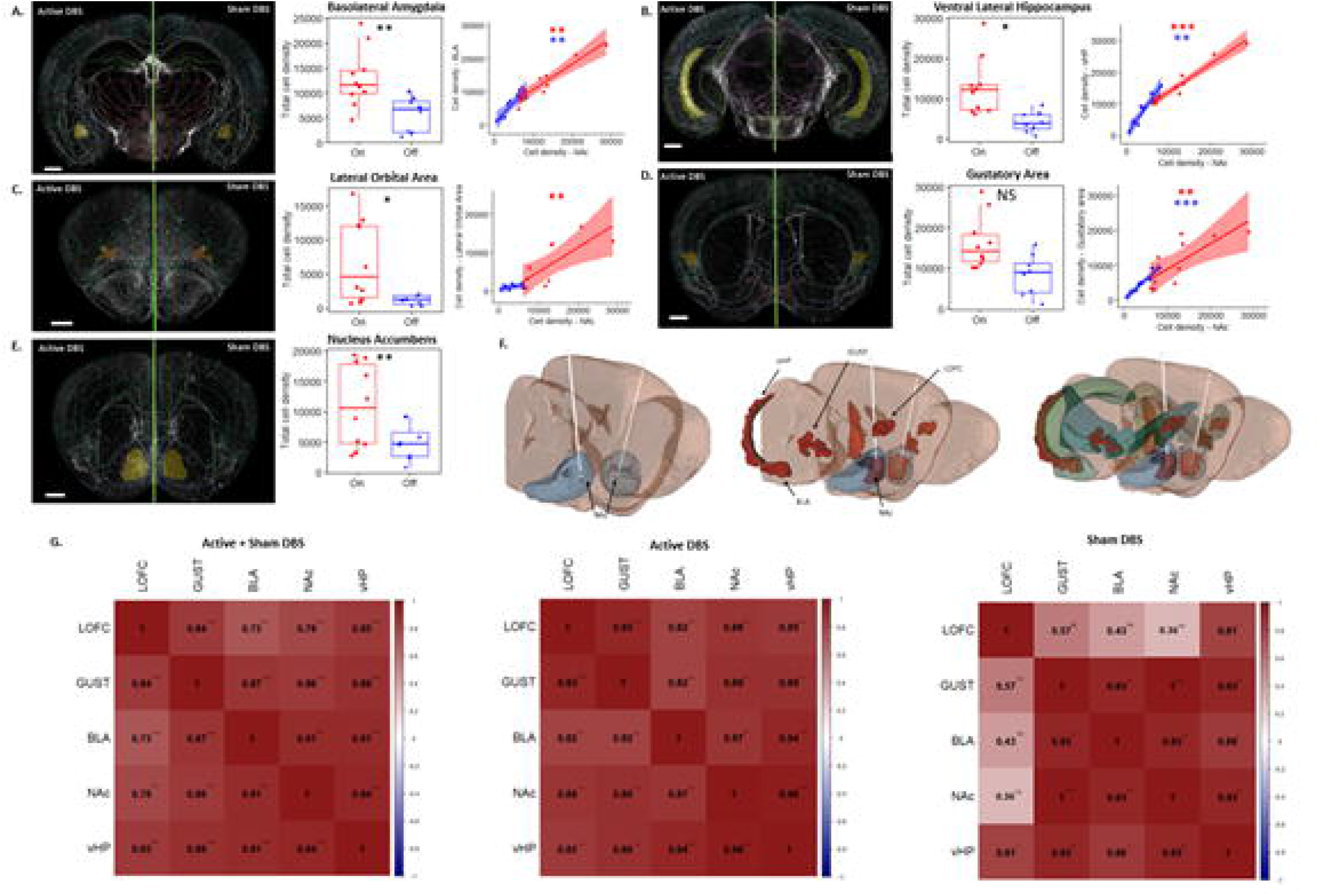
NAc DBS engages a preexisting network of mesolimbic sub-regions. **A**. Group-average Fos expression in BLA cluster for active versus sham DBS cohorts [left]. BLA cluster cell density is significantly greater in active DBS cohorts [center] and is very strongly correlated with NAc cluster cell density (*rho* = 0.93 / 0.87) for both sham and active DBS cohorts, respectively [right]. **B**. Group-average Fos expression in vHP neuronal cluster for active versus sham DBS cohorts [left]. vHP cluster cell density is significantly greater in active DBS cohorts [center] and is very strongly correlated with NAc cluster cell density (*rho* = 0.93 / 0.98) for both sham and active DBS cohorts, respectively [right]. **C**. Group-average Fos expression in LOFC neuronal cluster for active versus sham DBS cohorts [left]. Mean LOFC cell density is significantly greater in the active DBS group compared to sham DBS group [center]. The active DBS cohort is very strongly correlated with the NAc cluster cell density (*rho* = 0.88) while the sham DBS cohort is not (*rho* = 0.36) [right]. **D**. Group-average Fos expression in GUST neuronal cluster for active versus sham DBS cohorts [left]. Difference in GUST cell density between active and sham DBS is not significant [center], but both sham and active DBS are very strongly correlated with the NAc cluster cell density (*rho* = 1 / 0.89) [right]. **E**. Group-average FOS+ expression in NAc neuronal cluster for active versus sham DBS cohorts [left]. NAc cluster cell density is significantly greater in active DBS cohorts [right]. **F**. 3-dimensional mouse brain schematic of activated clusters. Leftmost displays bilateral electrodes with NAc shell targets (blue). Center displays all significant clusters (red) anatomically positioned – only right hemisphere clusters are labeled for clarity. Rightmost displays overlays of relevant regions in conjunction with clusters: hippocampus (light green), amygdala (light green), gustatory areas (dark green), and orbital areas (dark green). **G**. Correlations between all clusters. All clusters are strongly correlated with each other in the combined cohorts and the active DBS cohort. In the sham DBS cohort, some clusters’ correlations are weakened. Background voxel images in A-E are from group-average images of all hemispheres in active (left) and sham (right) DBS groups. Correlations were computed from Fos+ cell density in the respective clusters. Sham and active data are color-coded in red and blue, respectively. NS, P > 0.05; *P < 0.05; **P < 0.01; ***P < 0.001; BLA, basolateral amygdala; vHP, ventral hippocampus; LOFC, lateral orbitofrontal cortex; GUST, gustatory cortex.

To account for potential artifacts produced by normalization, smoothing, filtering and clustering of the statistical maps, we performed confirmatory analysis of single-cell Fos immunoreactivity at full resolution (3.5x3.5x3.5 um acquisition) in the original unfiltered brain volumes, within each of the 5 identified clusters.^39^ Within each cluster, we quantified the number of Fos+ cells per mm^3^ tissue (i.e., cell density) using Ilastik, an unbiased, supervised machine-learning-based method for pixel classification and cell counting. This approach bridges voxel-based intensity comparisons (i.e., permutation testing), which are a only a proxy for Fos expression, to a direct physiological measurement, the actual density of individual neuronal Fos expression, thus validating our previously identified clusters. We applied the previously generated transformation matrices to warp individual clusters from the standardized atlas space into the full-resolution native space of each sample. Active DBS cohorts had significantly greater average Fos+ cell densities in the BLA, vHP, LOFC, and NAc clusters (n = 36; BLA: W = 9; p = 0.004 | vHP: W = 13; p = 0.016 | LOFC: W = 12; p = 0.025 | NAc: W = 5; p = 0.001) compared to those of the sham cohorts (Fig. 2A, 2B, 2C and 2E). In the GUST, the difference between average total cell densities of the active and sham cohorts did not reach significance (GUST: W = 19; p = 0.068) (Fig. 2D). Fos+ cell density did not significantly differ between male and female mice for any cluster (Supplemental Fig. 1). To ensure that the results were not dependent on a specific cell counting algorithm, this cluster validation was performed with different cell counting algorithms (e.g., Ilastik object classification and Fiji 3D object counter; see “Cell Detection” in Methods for details) and cell densities within clusters were not significantly different across methods (Supplemental Fig. 2).

NAc DBS is associated with activation of distinct and noncontiguous clusters of Fos+ neurons across the BLA, vHP, and LOFC, suggesting a functionally connected network. We reasoned that correlated Fos+ density may reflect anatomical and functional connectivity in both active and sham DBS cohorts. Therefore, we examined covariance of Fos+ expression in pairs of clusters using data sham-treated and active DBS cohorts, both as separate cross-correlation analyses and in aggregate (N=36 hemispheres), to generate predictions about direct functional coupling (Fig. 2G). We found strong and statistically significant correlated Fos+ cell densities for every pair of clusters in the combined set of sham and active DBS cohorts (Fig. 2A-D, G *left*). Similar correlations were observed within active DBS cohort (Fig. 2G *center*). The strongest correlations with NAc Fos+ cell density in the active DBS cohort were observed with the BLA/vHP/LOFC clusters (*rho* = 0.87, 0.98, and 0.88, respectively; Fig. 2A-C, G *center*). In the GUST cluster, Fos+ cell density in the active DBS cohort was not significantly different from sham DBS, however the correlation of NAc and GUST Fos+ cell densities was nonetheless strong, suggesting anatomical connectivity (*rho* = 0.89; Fig. 2D, G *center*). Interestingly, in the sham DBS cohort, NAc Fos+ cell density remained significantly correlated with all but one cluster, the LOFC. Despite significantly higher Fos+ cell density in active DBS compared to sham in LOFC, correlation with NAc Fos+ density was weak and nonsignificant (*rho* = 0.36) in the sham cohort. Correlations of Fos+ density between LOFC and other clusters were also markedly weaker in the sham DBS cohort (GUST: *rho* = 0.57; BLA: *rho* = 0.43; vHP: *rho* = 0.81; Fig. 2G r*ight*) despite strongly correlated Fos+ density in the active DBS cohort. These results imply there is an intrinsic set of specific mesocorticolimbic structures that are connected at baseline and are dynamically reconfigured during NAc stimulation.

### Cross-species similarities in NAc DBS network engagement

Our brain-wide imaging of Fos expression in mice strongly suggests that NAc DBS engages a discrete functional network of brain regions. We next asked whether this NAc network is translatable to the human brain. One single subject with medically intractable epilepsy underwent stereo-electroencephalography (stereo-EEG) for seizure localization and was implanted with depth electrodes in a variety of brain regions for clinically indicated seizure mapping as previously reported.^57^ Among these electrodes, one trajectory traversed the NAc, which we used to interrogate projections between the NAc and mesocorticolimbic structures potentially engaged by NAc DBS, including hippocampus, amygdala, and insula which are analogous to the murine clusters that we identified previously. We used a highly novel and innovative algorithm to cluster the raw response shapes into basis profile curves (BPC), which are canonical electrophysiological motifs that uniquely characterize and group the different stimulation-evoked potentials measured in a specific recording site into clusters (Fig. 3A).^35,42^ This algorithm is a flexible approach to address the limitation of precursor methods where evoked potentials must fit a stereotyped shape to be quantified for connectivity strength.

**Figure 3.**
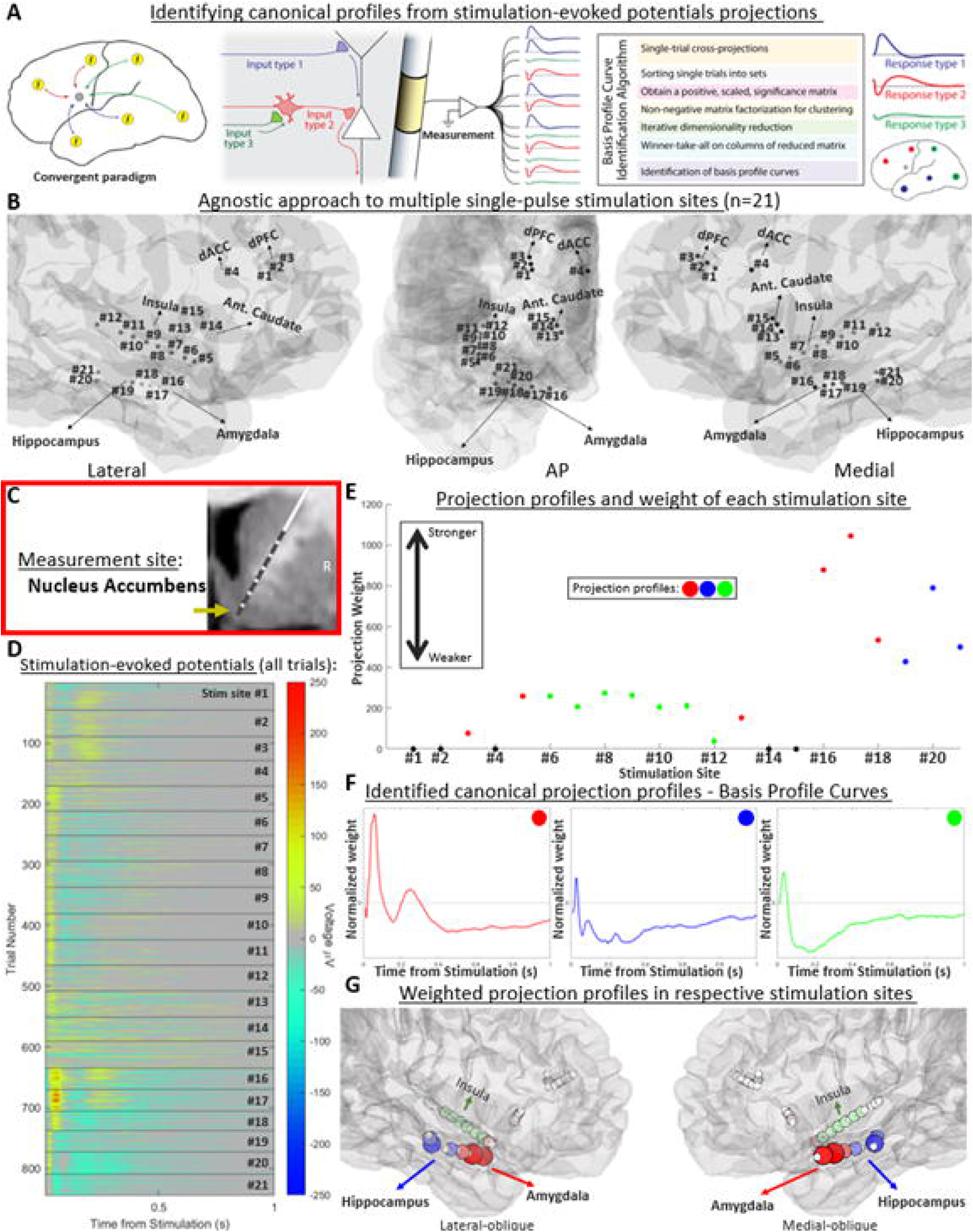
Map of NAc network engagement based on evoked potentials and the basis profile curve identification. **A**. Overview of method used to identify canonical profiles from stimulation-evoked potentials projections: Basis Profile Curves. **B**. Stimulation sites were overlaid onto their 3D position in the subject’s brain in their relative position (spatial average between both contacts used in bipolar stimulation of each site). **C**. Human NAc recording site. NAc field potentials were measured in a single electrode contact (gold arrow). **D**. Evoked potential matrix including all non-artefactual trials (n=844) across a total of 21 stimulation sites with multiple trials (average ∼40 trials/site, see Supplemental Material for traces resulting from each stimulation site). The observed exponential decay of the signal over time was secondary to an exponential decay weighting function applied to enhance focus on earlier signal changes as described in Huang et al, 2023. **E**. Classification of evoked potentials elicited by each stimulation site by BPC and signal magnitude. Each stimulation site is designated by its projection weight (group-averaged signal to noise ratio in relation to respective BPC) and color-coded by the underlying response motif (i.e., BPC). Stimulation pairs with similar anatomical localization often shared the same BPCs, as seen by the insula (green), amygdala (green), and hippocampus (red) groupings. **F**. BPCs calculated from evoked potential recordings. Each BPC is ∼1 second in length, starting from the cessation of stimulation. **G**. Anatomical location of BPCs. Every categorized projection was overlaid onto their 3D position in the subject’s brain as colored dots. Colors correspond to separate BPC motifs. Size and color intensity correspond to the strength of the projection weight (as quantified in D). White dots are the positions of the physical electrode contacts used in bipolar stimulation of each site.

We obtained direct recordings of human NAc activity in response to multiple trials of single-pulse bipolar stimulation for a total of 21 stimulation sites (Fig. 3B and Supplementary Figures for anatomical images of each stimulation site location). This convergent stimulation (Fig. 3A *left*) paradigm focused on the measurements (recordings) from a single site (NAc, Fig. 3C) and thus from a constant cytoarchitecture. When stimulating distant sites, different response shapes may thus reflect a variety of synaptic input configurations (Fig. 3A).^35,43^ After inspection and removal of trials containing artifacts (details in *Online Methods*), stimulation-evoked potentials recorded from the NAc in multiple trials (n = 844) of stimulation across all stimulation sites (average ∼40 trials per site) were selected for further analysis (Fig. 3D and Supplementary Figures for averaged traces elicited by each stimulation site). The BPC algorithm was applied to calculate the projection weight of each of the stimulation sites and assign their respective projection profile according to the BPC that best characterized the NAc evoked potentials (if any) measured in response to stimulation of that interconnected site (Fig. 3E). Three distinct BPCs were identified in the human NAc (Fig. 3F). The first BPC (red) waveform had a large, fast positive deflection observed within the initial 100 milliseconds, followed by a smaller and slower positive deflection observed within 300 milliseconds and a slow negative phase. The second (blue) and third (green) BPCs had small, fast positive deflections within 100 milliseconds followed by a slow and accentuated negative deflection within 200 milliseconds and a slower return to baseline in the second than the third BPC. The unique profiles of each BPC cluster imply differences in projection architecture. Evoked potentials classified with strong weights of the same BPCs cluster primarily corresponded to the stimulation sites in the hippocampus (blue, stimulation sites #19-21) and amygdala (red, #16-17), echoing prior findings in mice, as well as the insula (green, #6-12) (Fig. 3G), which at least in part is analogous to the murine gustatory areas. The fact that neuronal engagement mapped through mouse brain clearing in those regions could be reproduced with human single-pulse evoked potential experiments strongly implies a conserved network across species.

## Discussion

In the present study, we sought to elucidate the underlying network engagement of NAc DBS through novel, multimodal techniques across species.^9,10^ Using an unbiased, whole-brain imaging approach, NAc DBS was associate with an increased Fos expression in a variety of key mesolimbic structures. Furthermore, positive correlations in cell density were not only identified between these structures in the active DBS cohort, but were largely present in the sham cohort, representing an intrinsic functional network. Lastly, human single-pulse evoked potential (SPEP) recordings from the NAc generated unique basis profile curves that revealed strong similarity to the intrinsic functional network of NAc modulation seen in mice.

Current investigations of DBS circuitry and its network mechanisms rely heavily on functional imaging methods.^44,45^ These techniques can detect *in vivo* changes in the brain with millimetric spatial and reasonable temporal resolution during DBS. Indeed, our permutation analysis outputs have notable overlap with blood oxygen level dependent (BOLD) and cerebral blood volume (CBV) fMRI maps, reinforcing the presence of brain-wide modulation during NAc DBS.^44,45^ However, the discrepancies with those previously reported maps (i.e., activation of prefrontal cortex, ventral pallidum, etc.) may raise questions regarding reproducibility of NAc DBS effects as there is no standardized stereotactic target and non-selective nature of macrostimulation with the relatively large electrodes used in clinical settings.^57^ While functional imaging utilizes an indirect measurement of neuronal activity (e.g., blood oxygen level or blood volume), our methodology is very likely to be a more direct assessment of physiologically relevant processes, as Fos is translated in approximate proportion to the degree of neuronal firing.^46^ This fundamental difference in measurement of activation leads to differences in mapping, which has been evident in rats who received an identical intervention and were found to exhibit different fMRI and Fos activation maps.^47^ For instance, increased inhibitory synaptic activity could lead to a hemodynamic response but not an increase in Fos expression. Differences in experimental design also contribute to discrepancies in outcome. Rodent fMRI experiments have better temporal resolution compared to Fos expression and allow for repeated trials with the same subject, however live rodent imaging typically requires anesthesia, the time scale for stimulation is limited to seconds, and fMRI data has a limited signal-to-noise ratio.^44,45^ Comparatively, our study allows mice to freely roam with delivery of a much longer duration stimulation, though immediate sacrifice is necessary to measure Fos expression. Finally, we leverage in-human SPEPs, which have recently emerged as a powerful technique for studying regional interactions through the application of stimulation pulses at a specified site and measurement at another; SPEP refers to specifically to responses to short pulses separated by a brief interval of time (e.g., 2 seconds). Notwithstanding the high spatial and temporal resolution, the technique is limited by the number of electrode contacts and sampling bias.

Enhanced connectivity between NAc connected regions primarily during active NAc stimulation has previously been reported in fMRI studies with rodent and swine models.^44,45^ In the present work, very strong correlations of activated neuron density were found not only for the active DBS cohort, but also, perhaps not surprisingly, for the sham cohort. The existence of robust correlations pre-stimulation suggests that an intrinsic functional network may be present between the BLA-vHP-GUST-NAc clusters.

Our findings suggest that NAc DBS leads to increased neuronal activation across linked clusters, but still retaining roughly equivalent correlations between most clusters. This preservation indicates similar transmission mechanisms and projections are likely responsible, as opposed to the recruitment of new fibers outside an intrinsic network. Such a phenomenon would be physiologically consistent with axonal activation, which has been shown to underlie long-distance effects of DBS.^48^ As therapeutic DBS is known to alter neural activation patterns, the coordinated activation of the BLA-vHP-GUST-NAc may be a reflection of an efficacious pattern given our utilization of clinically relevant parameters.^16,49,50^ On the other hand, LOFC neuron modulation by NAc DBS likely may occur through another mechanism given significant contribution of GABAergic interneurons in the NAc-OFC connection amongst others.^30,51^ Through presumed activation of interneurons that inhibit excitatory pyramidal neurons, reduction of overall OFC neuron firing has been observed.^52^ This complex orchestration of excitation and inhibition could be consistent with the observed insignificant change in Fos expression as the interneuron Fos expression could be offset by the suppression of pyramidal neurons.

Complementing our mouse NAc DBS mesocorticolimibic network mapping, BPCs were generated from rare NAc recordings of a human subject with depth electrodes traversing multiple mesocorticolimbic structures. Bidirectional stimulation could not be performed as the size of the implanted NAc electrode would result in nonspecific stimulation of both NAc and neighboring regions (e.g., ventral anterior limb of internal capsule, caudate, olfactory tuberculum, etc.), and the human intracranial electrophysiology setup requires bipolar stimulation via a pair of electrodes, which would not yield comparable anatomical specificity to recording from a single electrode placed in NAc. Moreover, the convergent approach employed here facilitates an assessment of the stimulated interconnections while controlling for underlying recording cytoarchitecture with a single the recording site. The spatial distribution of the different BPCs closely parallels that of the separate electrodes, suggesting unique interconnections with each brain region. Indeed, similar anatomical distributions of BPCs by region are seen in a recent study of human ventral temporal cortex SPEP recordings.^42^ The amygdala-NAc, hippocampal-NAc, and insula-NAc BPCs likely reflect the differing macro (e.g., cell types and laminar structure) and microscale (e.g., ion flow) nature of these different connections. The information carried by SPEP waveforms has been thoroughly explored, from afferent excitatory postsynaptic potential coherence to underlying brain rhythms.^43,53^ However, without a convergent paradigm as utilized with BPCs, interpretation and translatability is limited by differing input and output architecture. For instance, the commonly accepted N1/N2 response was not sufficient to describe the complex inputs from many different cortical regions to the parahippocampal gyrus or ventral temporal cortex.^35,42^ As tools for reimaging motif dynamics, BPCs may be able to provide novel insights into network engagement. The translatability of the stimulation clusters from murine to human not only reinforces their physiological validity, but also indicates the potential for using brain clearing techniques to elucidate and study stimulation networks underlying clinically-relevant neuromodulatory approaches. Stimulation maps may elucidate and enable further characterization of neurons of downstream targets in combination with labeling techniques. The typical and canonical nature of BPCs facilitates inferences about underlying architectures and processes - a library of BPCs would be a powerful addition for neural mapping. Consensus BPCs across individuals have recently been demonstrated in proof-of-concept work for ventral temporal cortex recordings.^42^

Our objective was to better understand network engagement by a promising DBS target for neuropsychiatric disorders. It needs to be acknowledged that an unavoidable difference exists in the technical implementation of human intracranial electrophysiology and and mouse DBS. Consequently, a sampling bias may occur in the former given limitations in implant sites (e.g., no human LOFC electrode) while 3D imaging is unbiased in the latter. Furthermore, while the convergent paradigm allowed pinpointing the SPEP to a single NAc electrode, for instance, the size of human bipolar stimulation fields limits specification of stimulated amygdala, hippocampal, and insular sub-regions. The overall impact of these differences would be difficult to quantify but the potential downside is acceptable given the scope and purpose of this work. Another limitation of this study was the scope of the experimental design with a single set of stimulation settings in a healthy rodent model. While the Fos expression does not necessarily reflect only acute changes in activity, days or weeks of chronic stimulation may result in substantially different patterns of Fos expression.^15^ In addition, more robust markers of neuronal activation could be considered in the future. Another set of limitations to our work may result from the discrepancy between measured and actual network engagement. Atlas registration of mouse brains process may be imprecise, particularly for smaller brain regions. Additionally, as DBS increases Fos expression, activated regions with dense Fos expression at baseline may be less apparent due to a ceiling effect.

In the present study, we have elucidated a network of brain regions activated by NAc DBS through an unbiased whole-brain clearing and cell detection pipeline. We have also used a convergent approach in a human subject to characterize canonical stimulation motifs in the NAc during single-pulse stimulation of these regions. Our findings not only illustrate a novel methodology for uncovering mechanisms of focal interventions modulating brain networks, but it also expands our understanding of NAc DBS network engagement, which may guide future clinical trials of NAc-network based stimulation.

## Acknowledgments

We thank the participants who took part in the study. We thank Drs. Johnathon J. Parker and Rajat Shivacharan, and for their support acquiring the single-pulse stimulation data in the Epilepsy Monitoring Unit. We are grateful to Dr. Josef Parvizi for his invaluable support that enabled the electrophysiology data acquisition in the Epilepsy Monitoring Unit. This work was supported by Stanford University and University of Pennsylvania Neurosurgery start-up funds, the John A. Blume Foundation, the William Randolph Hearst Foundation, the Foundation for OCD Research (New Venture Fund), the AE Foundation and the National Institutes of Health (RO1 MH124760).

## Author Contributions

Conceptualization: A.Y.F., D.A.N.B., C.H.H., and B.D.H.; Methodology: A.Y.F., D.A.N.B., C.H.H., D.R.R., K.J.M., and B.D.H.; Formal Analysis: A.Y.F., D.A.N.B., A.B.C., D.R.R., and J.S.S.; Investigation: A.Y.F., J.S.S., and D.A.N.B.; Resources: R.C.M., C.H.H., and B.D.H.; Writing – Original Draft: A.Y.F. and D.A.N.B.; Writing – Review & Editing: A.Y.F., D.A.N.B., C.H.H., D.R.R., H.H., D.H., K.J.M., and B.D.H.; Visualization: A.Y.F. and D.A.N.B.; Supervision: R.C.M., C.H.H., and B.D.H.

## STAR Methods

### Experimental model and study participant details

For the present work, we utilized 8-week-old TRAP2:Ai14 mice (N = 18) with double homozygous genotype confirmed at 2 weeks age with tail biopsies. This mouse line is a recently developed tool that allows for capture-and-control of neural circuits defined by their activity within a temporal window of 4-6 hours, triggered by administration of 4-OHT. While TRAP2 mice have been used by multiple groups to interrogate the role of functionally defined circuits, TRAP2 reporter expression (via Ai14) does not uniformly represent Fos expression across the brain.^54–56^ By examining Fos in mice that also express tdT, we are able to establish which identified circuits (by Fos mapping) may be amenable to manipulation using the TRAP2 mouse line, facilitating future work by our and other groups.

Equal numbers of male (n = 9) and female (n = 9) mice were utilized. All mice were individually housed in plastic cages with disposable bedding on a standard light cycle with *ad libitum* access to standard chow and water. All procedures followed the National Institutes of Health Guide for the Care and Use of Laboratory Animals and received approval from the Stanford University Administrative Panel on Laboratory Animal Care (#32691).

### Method details

#### Animal DBS

All mice were implanted with bipolar electrodes (FHC, Inc., Bowdoin, ME, USA) in the NAc shell bilaterally. They were anesthetized with isoflurane (5% induction, 1% maintenance) and mounted to a stereotaxic frame (David Kopf Instruments, Tujunga, CA, USA). The NAc shell region is targeted with the following coordinates relative to bregma: +1.35 mm anterior-posterior, -4.25 mm dorsal-ventral, and +/- 0.6 mm lateral-medial (Fig 1A). Electrodes are inserted at a 10-degree angle relative to the orthogonal direction to facilitate bilateral placement. Pain was managed with meloxicam (0.5 mg/mL) and all animals were monitored until recovery from anesthesia. Experiments started 1-week post-electrode implantation to allow for healing.

Mice are divided into 2 stimulation cohorts: active and sham. Both active (N = 10 / 10) and sham (N = 8 / 8) cohorts had equal numbers of male and female mice (Fig 1B). Following recovery from electrode implantation, mice are first transferred in their home cages into the experimental chambers where wires are attached without any electrical stimulation to facilitate habituation. After 1 hour, stimulation delivered as continuous biphasic pulses (130 Hz, 0.1 mA, 90 µs) was started for the active DBS cohort. One hour after DBS initiation, mice in both cohorts received intraperitoneal injections of 4-hydroxytamoxifen (4-OHT) at the lower right quadrant of the abdomen. DBS was terminated for the active DBS cohort 3 hours after the 4-OHT injection. All mice remained in their experimental chambers with attached wires for an additional 3 hours to minimize extraneous neural stimulation and possible Fos expression. Mice were then detached and removed from the experimental chambers. Overall, mice in the active DBS cohort received a total of 4 hours of stimulation. No electrical stimulation was delivered to the sham cohort at any point.

At the second stimulation experiment 1 week later, mice were again given 1 hour to habituate to the experimental chambers and attached wires. Afterwards, stimulation identical to that mentioned above was initialized for the active DBS cohort. No stimulation was started in the sham cohort. Termination of DBS occurred after 1 hour, at which point all mice were immediately detached.

#### iDISCO brain-clearing 3D histology

Immediately following the second stimulation, all mice were killed by trans-cardial perfusion. Following anesthetization with 5% isoflurane, mice were trans-cardially perfused with 20 mL 1x phosphate-buffered saline (PBS). Next, 20 mL of ice-cold 1x PBS / 4% paraformaldehyde (PFA) were perfused. Electrodes were first carefully removed. Whole brains were extracted from the crania and post-fixed for 24 hours in 1x PBS / 4 % PFA at 4°C. All solutions were freshly prepared from stock.

##### Pretreatment

Fixed brains are removed from 4°C and allowed to warm to room temperature. They are moved to new tubes (5 mL Eppendorf) and rinsed with 1x PBS with nutation for 30 minutes, repeated 3 times. Samples are returned to 4°C in 1x PBS overnight. On day 2, samples are moved into new tubes and progressively dehydrated with solutions of increasing concentrations of methanol in PBS (20%, 40%, 60%, 80%, and 100%) for 1 hour each with shaking. An additional wash with 100% methanol for 1 hour is performed. All samples are chilled to 4°C for 20 minutes before an overnight incubation in a solution of 66% dichloromethane (DCM) / 33% methanol with nutation at room temperature. On day 3, samples are washed twice with 100% methanol for 1 hour at room temperature before being transferred to new tubes. Freshly prepared and chilled 5% hydrogen peroxide in methanol is used to bleach the samples overnight at 4°C without shaking. On day 4, samples are moved into new tubes and rehydrated with solutions of decreasing concentrations of methanol in PBS (80%, 60%, 40%, 20%, and 0%) for 1 hour each with shaking. Samples are again moved into new tubes and washed for 1 hour in PTx.2 (100 mL PBS 10x + 2 mL TritonX-100). After transferring into new tubes, samples are incubated in a freshly prepared permeabilization solution (400 mL PTx.2, 11.5 g glycine, 100 mL dimethyl sulfoxide (DMSO)) at 37°C for 48 hours with nutation. At 24 hours, samples are transferred into new tubes with fresh permeabilization solution before returning to the incubator.

##### Immunolabeling

After moving into new tubes, samples are incubated in freshly prepared blocking solutions (75.6 mL PTx.2 + 5.4 mL donkey serum (DS) + 3 mL DMSO) at 37°C for 48 hours with rotation (Fig 1C). Samples are then moved into new tubes and incubated with the primary antibodies – rabbit anti-Fos (1:500; Synaptic systems, Goettingen, Germany, 226-008) and chicken anti-RFP (1:500; Novus Biologicals, Littleton, CO, USA, NBP1-97371) – diluted in freshly prepared solution of PTwH [100 mL PBS 10X, 2 mL Tween-20, 1 mL of 10 mg/mL Heparin stock solution] / 5% DMSO / 3% DS at 37°C for 10 days with nutation. At completion of primary incubation, samples are moved to new tubes, washed with PTwH for several hours, and left overnight nutating. Next, samples are incubated with the secondary antibodies – goat anti-chicken Alexa 568 (1:250; Thermo Fisher Scientific Inc., Waltham, MA, USA, A11041) and donkey anti-rabbit Alexa 647 (1:250; Thermo Fisher Scientific Inc., Waltham, MA, USA, A31573) – diluted in PTwH / 3% DS at 37°C for 10 days with nutation and limited light exposure. At completion of secondary incubation, samples are washed with PTwH with nutating for 48 hours.

##### Clearing

Dried samples are embedded in 1% low melting point agarose and dehydrated with solutions of increasing concentrations of methanol in PBS (20%, 40%, 60%, 80%, and 100%) for 1 hour each with nutation. An additional wash with 100% methanol for 1 hour is performed after moving to new tubes.

Samples are covered and left to incubate overnight in 66% DCM / 33% methanol at room temperature. On the following day, samples are first washed with 100% DCM for 15 minutes twice with nutation at room temperature. Next, samples are moved to a new tube and incubated with dibenzyl ether with no agitation. Contact with air and light should be minimized for the samples.

### Light sheet Imaging and preprocessing

The multi-modal image registration and connectivity analysis (MIRACL) tool is an open-source, analysis pipeline that facilitates the various preprocessing and registration steps that result in seamless overlap between atlas and cleared sample data.^41^ The light sheet fluorescence microscopy (LSFM) atlas, which is an optimized digital mouse brain atlas specifically designed for light sheet fluorescence microscopy, is used in this project. After conversion from CZI to TIFF file format, image dimensions are cropped if not even values. Background subtracted images are generated through rolling ball subtraction. Rolling ball calculates and subtracts local background values for every pixel by averaging a large ball around the pixel, thereby hypothetically removing any spatial variations in background intensities; a rolling ball radius of 4 was used (Fig 1D).^57^ Next, images are down-sampled by a factor of 2 and converted to NIFTI format. The co-registration was performed using the MIRACL pipeline, as previously described, to register samples to a standard atlas, substituting an averaged template brain generated with iDISCO+ and LSFM in place of one obtained via serial two-photon microscopy, to better match our samples’ autofluorescence profile (Fig. 1E). This resulted in the transformation matrix that allows conversion of the LSFM atlas in template space into native space (Fig 1E).

### Cell detection

Ilastik is an unbiased, interactive segmentation program that was used to classify active cells.^37^ For “pixel classification”, a Random Forest classifier uses user annotations and selected pixel features to generate a binary probabilistic map. Raw data (TIFFs) were imported as stacks of 20 axial slices of X thickness. Representation of both ON (N = 3) and OFF (N = 3) cohorts was balanced for training the classifier. The following parameters were chosen for pixel features: Gaussian Smoothing (0.3 & 3.5), Laplacian of Gaussian (3.5 & 5.0), Gaussian Gradient Magnitude (3.5), and Hessian of Gaussian Eigenvalues (0.7 & 1.6). Individual Z-stacks are then annotated as ‘cell’ or ‘non-cell’ regions. The “object classification” workflow builds on pixel classification with additional training, ultimately designating sets of pixels as belonging to single entities. Hysteresis thresholding was utilized with x, y, z.

Fiji is an open-source multifunctional image processing software.^58^ Its 3D object counter is a plugin that quantifies 3D objects within a stack. For parameters, the size filter minimum and maximum are x and y, respectively, and edge objects were excluded. This cell detection method inputs the Ilastik pixel classification outputs, which is a binarized map of background and classified cells.

Given the heavy computational burden of performing cell detection on full samples, both methods were instead run on volumes of the cluster. To generate these volumes, the length, width, and height of full-resolution clusters were obtained through the statistics tool of the FMRIB Software Library (FSL). After cropping out this specific volume from the entire brain, a binary cluster mask was applied to this new volume. This process was performed on raw data or the Ilastik pixel classification output for the Ilastik object classification and Fiji 3D object counter, respectively. Ilastik object classification was the primary cell detection method given the more comprehensive classification methodology and its non-inferiority to the Fiji 3D cell counter.

## Statistical Analyses

### Human Stimulation-Evoked Potentials

#### Stereo-EEG surgery and electrode localization

The participant was a patient with refractory epilepsy whose pattern of epileptiform activity warranted invasive brain mapping as part of additional standard-of-care work-up. After a meeting between her treatment team, a consensus decision was made to implant the patient’s brain with intracranial electrodes in the suspected cortical regions such as the frontal lobe, the hypothalamus (through the nucleus accumbens), and the limbic lobe including medial temporal lobe, insula, anterior and anterior cingulate.^59^ The entire insula was evaluated with an anterior to posterior trajectory.

Stereo-EEG procedure with depth electrodes (0.86 mm in diameter; 2.29 mm height; 3–5 mm inter-electrode center-to-center distance made by *AdTech*). Pre-surgical MRI scan was co-registered to post-surgical CT scan for electrode visualization and localization as described previously (ref). Locations of depth electrodes utilized in single-pulse stimulation experiments were then examined by one rater with expertise in neuroanatomy and neuroimaging (D.A.N.B). All electrodes utilized in those experiments as well as a recording electrode in the NAc were selected for further evaluation and were labeled numerically. This localization was performed prior to signal processing

#### Single pulse evoked potentials acquisition and pre-processing

As previously described, we performed single-pulse stimulations at rest using an intracranial electrical waveform generator and switchbox (MS-120BK-EEG and PE-210AK, Nihon Kohden, Tokyo, Japan).^60,61^ Electrical stimulation was delivered through adjacent pairs of electrodes in biphasic pulses (6mA; 200 μs/phase, 49 trials) at a frequency of 0.5 Hz for a total of 120 seconds. We measured electrical potentials in the NAc in response to stimulation with a video EEG monitoring system using a sampling rate of 2000 Hz (version WEE-1200, Nihon Kohden). We analyzed the single-pulse stimulation data using custom Matlab scripts. We first applied a high-pass butterworth filter (1 Hz) to exclude slow varying effects and segmented evoked responses time series from recording channels were into 2500 ms epochs time-locked to stimulus onset (500 ms pre-stimulus and 2000 ms post-stimulus). Then, we re-referenced the data to the common average signal, excluding stimulated channels, channels with artifacts, and channels with large, evoked responses, as previously described.^35^ Finally, to exclude potential effects of pre-stimulus signal fluctuations, we applied a baseline correction by subtracting the average signal between 500 ms and 10 ms prior to stimulus onset. Single pulse evoked potentials (SPEP) were obtained from the time series of the NAc recordings as single pulse simulations were performed. An exponential decay weighting function with time constant, t, equal to 500 ms was applied to all trials to focus on earlier signal changes following stimulation as previously described.^42^

#### Basis profile curve identification

Given the limitations and ambiguity in interpreting SPEP measurements, a novel algorithm involving a convergent paradigm for generating basis profile curves (BPC) was utilized.^35^ BPCs are canonical summaries of similar evoked potential traces, which facilitates a higher-level quantification of regions under investigation. For the present study, the NAc (electrode location) was the single recording site with the following stimulation electrode contacts (Table / figure). Stimulation is delivered as biphasic pulses to all electrode pairs, with each trial unit-normalized and stored in a matrix 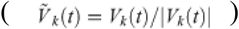. Self and cross-projections of single trials are performed 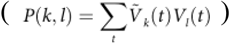 and sorted into different subgroup interaction sets (*S*_n,m_ = *P {k* ∈ *n*,*l* ∈ *m*}). Next, a scaled significance matrix (t-values from Sn,m) is constructed. Of note, negative t-values need to be set to zero in order to fulfill a non-negativity constraint. As the convergent approach eliminates the problem of source-localization, an individual cluster’s contribution cannot be both positive and negative because the underlying architecture of the single recording site is stable. Hence, if there are flips in voltage at a stimulation site, the constraint facilitates segregation into different clusters. To identify and categorize different CCEP clusters, non-negative matrix factorization (NNMF) is used to generate the decomposition: Ξ∼ M H. Ξ has dimensions of *N x M*. Dimensions for W and H are *N x Q* and *Q x M*, respectively. W and H are updated through a repetitive multiplication process. First, elements in H are scaled as:

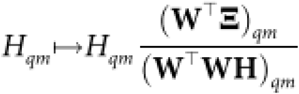

Second, rows of H are unit normalized and the columns of W are multiplied by the normalization factor:

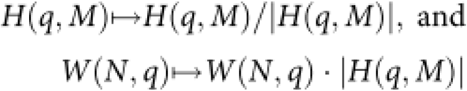

Third, elements of W are scaled:

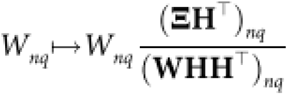

Lastly, error ( η|Ξ −WHΞ ^2^ ) is calculated. The process is repeated until the error reaches a set tlΔη/η<10−5.Given the highly degenerate output (off-diagonal elements of matrix **HH**^**T**^) of NNMF, dimensionality reduction is accomplished by iteratively reducing Q (inner component number) by 1 until z (maximum of the upper-half off-diagonal elements of **HH**^**T**^) < 0.5. This process limits the quantity of shared structures between different response motifs. A “winner-takes-all” approach is adopted for each column in **H** (non-maximal elements are set to zero) such that each subgroup element only belongs to a single component. For each *q* row in **H**, subgroup *n* is assigned to a cluster with thresholding if 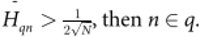 then *n* ∈ *q* . Lastly, a linear kernel principal component decomposition (PCA) is used to invert the decomposition and identify a representative basis curve. Individual trials can be generated with the BPC with a scalar and residual error. Specific details regarding methodology can be found in Miller et al. 2021.

##### Quantification and statistical analysis

Identification of significant clusters is accomplished through nonparametric permutation thresholding with the *randomise* function of FSL. The data, as a 4D (time-series of 3D images) NIFTI file, are modeled in a generalized linear model (GLM) with design (to designate independent variables) and contrast (to compare variable outputs) matrices (Fig 4F). Threshold-free cluster enhancement, a methodology that enhances cluster-like structures without preset threshold, was used for the output. The null distribution was generated with 5000 permutations. Given the resolution of the input images and consideration of physiological feasibility, a minimal volume of 2500 voxels were set a priori. The output was further refined with a smoothing sigma of 50 um and a threshold of p < 0.01, yielding the final clusters.

Cluster cell densities are defined as the number of active cells in the cluster divided by the cluster volume. Total cluster cell densities are the summation of both left and right cluster cell densities. Statistical analyses were performed using R Studio version 3.6.3. Wilcoxon tests were performed to compare mean total cell densities (active cell count / cluster volume) between ON and OFF cohort samples. Grubbs’ test was performed to detect outliers (normality assumption was evaluated with the Shapiro-Wilk test). Spearman’s rank-order correlations were used to calculate the association between different neuronal clusters. Statistical significance was determined using alpha = 0.05.

## Supplemental Information

Document S1: Figures S1-S5

## References

1. Kessler, R.C., Aguilar-Gaxiola, S., Alonso, J., Chatterji, S., Lee, S., Ormel, J., Üstün, T.B., and Wang, P.S. (2009). The global burden of mental disorders: An update from the WHO World Mental Health (WMH) Surveys. Epidemiol. Psichiatr. Soc. 18, 23. 10.1017/S1121189X00001421.

2. Howes, O.D., Thase, M.E., and Pillinger, T. (2022). Treatment resistance in psychiatry: state of the art and new directions. Mol. Psychiatry 27, 58. 10.1038/S41380-021-01200-3.

3. Russo, S.J., and Nestler, E.J. (2013). The Brain Reward Circuitry in Mood Disorders. Nat. Rev. Neurosci. 14, 609–625. 10.1038/NRN3381.

4. Koob, G.F., and Volkow, N.D. (2016). Neurobiology of addiction: a neurocircuitry analysis. The lancet. Psychiatry 3, 760. 10.1016/S2215-0366(16)00104-8.

5. Baxter, L.R., Schwartz, J.M., Bergman, K.S., Szuba, M.P., Guze, B.H., Mazziotta, J.C., Alazraki, A., Selin, C.E., Ferng, H.K., Munford, P., et al. (1992). Caudate Glucose Metabolic Rate Changes With Both Drug and Behavior Therapy for Obsessive-Compulsive Disorder. Arch. Gen. Psychiatry 49, 681–689. 10.1001/ARCHPSYC.1992.01820090009002.

6. Everitt, B.J., and Wolf, M.E. (2002). Psychomotor Stimulant Addiction: A Neural Systems Perspective. J. Neurosci. 22, 3312. 10.1523/JNEUROSCI.22-09-03312.2002.

7. Lüscher, C., Robbins, T.W., and Everitt, B.J. (2020). The transition to compulsion in addiction. Nat. Rev. Neurosci. 21, 247. 10.1038/S41583-020-0289-Z.

8. Klawonn, A.M., and Malenka, R.C. (2018). Nucleus Accumbens Modulation in Reward and Aversion. Cold Spring Harb. Symp. Quant. Biol. 83, 119. 10.1101/SQB.2018.83.037457.

9. Haber, S.N., and McFarland, N.R. (1999). The Concept of the Ventral Striatum in Nonhuman Primates. Ann. N. Y. Acad. Sci. 877, 33–48. 10.1111/J.1749-6632.1999.TB09259.X.

10. Haber, S.N., Lynd, E., Klein, C., and Groenewegen, H.J. (1990). Topographic organization of the ventral striatal efferent projections in the rhesus monkey: An anterograde tracing study. J. Comp. Neurol. 293, 282–298. 10.1002/CNE.902930210.

11. Zahm, D.S., and Heimer, L. (1993). Specificity in the efferent projections of the nucleus accumbens in the rat: Comparison of the rostral pole projection patterns with those of the core and shell. J. Comp. Neurol. 327, 220–232. 10.1002/CNE.903270205.

12. French, S.J., and Totterdell, S. (2003). Individual nucleus accumbens-projection neurons receive both basolateral amygdala and ventral subicular afferents in rats. Neuroscience 119, 19–31. 10.1016/S0306-4522(03)00150-7.

13. French, S.J., and Totterdell, S. (2002). Hippocampal and prefrontal cortical inputs monosynaptically converge with individual projection neurons of the nucleus accumbens. J. Comp. Neurol. 446, 151–165. 10.1002/CNE.10191.

14. Al-Hasani, R., Gowrishankar, R., Schmitz, G.P., Pedersen, C.E., Marcus, D.J., Shirley, S.E., Hobbs, T.E., Elerding, A.J., Renaud, S.J., Jing, M., et al. (2021). Ventral tegmental area GABAergic inhibition of ventral accumbens shell cholinergic interneurons promotes reward reinforcement. Nat. Neurosci. 24, 1414. 10.1038/S41593-021-00898-2.

15. Wu, H., Kakusa, B., Neuner, S., Christoffel, D.J., Heifets, B.D., Malenka, R.C., and Halpern, C.H. (2022). Local accumbens in vivo imaging during deep brain stimulation reveals a strategy-dependent amelioration of hedonic feeding. Proc. Natl. Acad. Sci. U. S. A. 119, e2109269118. 10.1073/PNAS.2109269118/SUPPL_FILE/PNAS.2109269118.SAPP.PDF.

16. Halpern, C.H., Tekriwal, A., Santollo, J., Keating, J.G., Wolf, J.A., Daniels, D., and Bale, T.L. (2013). Amelioration of Binge Eating by Nucleus Accumbens Shell Deep Brain Stimulation in Mice Involves D2 Receptor Modulation. 10.1523/JNEUROSCI.3237-12.2013.

17. Barbosa, D.A.N., Kuijper, F.M., Duda, J., Wang, A.R., Cartmell, S.C.D., Saluja, S., Cunningham, T., Shivacharan, R.S., Bhati, M.T., Safer, D.L., et al. (2022). Aberrant impulse control circuitry in obesity. Mol. Psychiatry 27, 3374. 10.1038/S41380-022-01640-5.

18. Wang, A.R., Kuijper, F.M., Barbosa, D.A.N., Hagan, K.E., Lee, E., Tong, E., Choi, E.Y., McNab, J.A., Bohon, C., and Halpern, C.H. (2023). Human habit neural circuitry may be perturbed in eating disorders. Sci. Transl. Med. 15, eabo4919. 10.1126/SCITRANSLMED.ABO4919/SUPPL_FILE/SCITRANSLMED.ABO4919_MDAR_REPRODUCIBILITY_CHECKLIST.PDF.

19. Follett, K.A., Weaver, F.M., Stern, M., Hur, K., Harris, C.L., Luo, P., Marks, W.J., Rothlind, J., Sagher, O., Moy, C., et al. (2010). Pallidal versus Subthalamic Deep-Brain Stimulation for Parkinson’s Disease. N. Engl. J. Med. 362, 2077–2091. 10.1056/NEJMOA0907083/SUPPL_FILE/NEJM_FOLLETT_2077SA1.PDF.

20. Schuepbach, W.M.M., Rau, J., Knudsen, K., Volkmann, J., Krack, P., Timmermann, L., Hälbig, T.D., Hesekamp, H., Navarro, S.M., Meier, N., et al. (2013). Neurostimulation for Parkinson’s Disease with Early Motor Complications. 10.1056/NEJMoa1205158 368, 610–622. 10.1056/NEJMOA1205158.

21. DeHemptinne, C., Swann, N.C., Ostrem, J.L., Ryapolova-Webb, E.S., San Luciano, M., Galifianakis, N.B., and Starr, P.A. (2015). Therapeutic deep brain stimulation reduces cortical phase-amplitude coupling in Parkinson’s disease. Nat. Neurosci. 18, 779. 10.1038/NN.3997.

22. Shivacharan, R.S., Rolle, C.E., Barbosa, D.A.N., Cunningham, T.N., Feng, A., Johnson, N.D., Safer, D.L., Bohon, C., Keller, C., Buch, V.P., et al. (2022). Pilot study of responsive nucleus accumbens deep brain stimulation for loss-of-control eating. Nat. Med. 2022 289 28, 1791–1796. 10.1038/s41591-022-01941-w.

23. Scangos, K.W., Khambhati, A.N., Daly, P.M., Makhoul, G.S., Sugrue, L.P., Zamanian, H., Liu, T.X., Rao, V.R., Sellers, K.K., Dawes, H.E., et al. (2021). Closed-loop neuromodulation in an individual with treatment-resistant depression. Nat. Med. 27, 1696–1700. 10.1038/S41591-021-01480-W.

24. Davidson, B., Giacobbe, P., George, T.P., Nestor, S.M., Rabin, J.S., Goubran, M., Nyman, A.J., Baskaran, A., Meng, Y., Pople, C.B., et al. (2022). Deep brain stimulation of the nucleus accumbens in the treatment of severe alcohol use disorder: a phase I pilot trial. 10.1038/s41380-022-01677-6.

25. Kuijper, F.M., Mahajan, U.V., Ku, S., Barbosa, D.A.N., Alessi, S.M., Stein, S.C., Kampman, K.M., Bentzley, B.S., and Halpern, C.H. (2022). Deep Brain Stimulation Compared With Contingency Management for the Treatment of Cocaine Use Disorders: A Threshold and Cost-Effectiveness Analysis. Neuromodulation 25, 253–262. 10.1111/ner.13410.

26. Visser-Vandewalle, V., Andrade, P., Mosley, P.E., Greenberg, B.D., Schuurman, R., McLaughlin, N.C., Voon, V., Krack, P., Foote, K.D., Mayberg, H.S., et al. (2022). Deep brain stimulation for obsessive–compulsive disorder: a crisis of access. Nat. Med. 2022 288 28, 1529–1532. 10.1038/s41591-022-01879-z.

27. Guehl, D., Benazzouz, A., Aouizerate, B., Cuny, E., Rotgé, J.Y., Rougier, A., Tignol, J., Bioulac, B., and Burbaud, P. (2008). Neuronal Correlates of Obsessions in the Caudate Nucleus. Biol. Psychiatry 63, 557–562. 10.1016/j.biopsych.2007.06.023.

28. Denys, D., Mantione, M., Figee, M., Van DenMunckhof, P., Koerselman, F., Westenberg, H., Bosch, A., and Schuurman, R. (2010). Deep Brain Stimulation of the Nucleus Accumbens for Treatment-Refractory Obsessive-Compulsive Disorder. Arch. Gen. Psychiatry 67, 1061–1068. 10.1001/ARCHGENPSYCHIATRY.2010.122.

29. Bewernick, B.H., Kayser, S., Sturm, V., and Schlaepfer, T.E. (2012). Long-term effects of nucleus accumbens deep brain stimulation in treatment-resistant depression: evidence for sustained efficacy. Neuropsychopharmacology 37, 1975–1985. 10.1038/NPP.2012.44.

30. Vassoler, F.M., White, S.L., Hopkins, T.J., Guercio, L.A., Espallergues, J., Berton, O., Schmidt, H.D., and Christopher Pierce, R. (2013). Deep brain stimulation of the nucleus accumbens shell attenuates cocaine reinstatement through local and antidromic activation. J. Neurosci. 33, 14446–14454. 10.1523/JNEUROSCI.4804-12.2013.

31. Schmuckermair, C., Gaburro, S., Sah, A., Landgraf, R., Sartori, S.B., and Singewald, N. (2013). Behavioral and neurobiological effects of deep brain stimulation in a mouse model of high anxiety- and depression-like behavior. Neuropsychopharmacology 38, 1234–1244. 10.1038/NPP.2013.21.

32. Ho, A.L., Feng, A.Y., Barbosa, D.A.N., Wu, H., Smith, M.L., Malenka, R.C., Tass, P.A., and Halpern, C.H. (2021). Accumbens coordinated reset stimulation in mice exhibits ameliorating aftereffects on binge alcohol drinking. Brain Stimul. 14, 330–334. 10.1016/j.brs.2021.01.015.

33. Denys, D., Mantione, M., Figee, M., Van DenMunckhof, P., Koerselman, F., Westenberg, H., Bosch, A., and Schuurman, R. (2010). Deep Brain Stimulation of the Nucleus Accumbens for Treatment-Refractory Obsessive-Compulsive Disorder. Arch. Gen. Psychiatry 67, 1061–1068. 10.1001/ARCHGENPSYCHIATRY.2010.122.

34. Ho, A.L., Salib, A.M.N., Pendharkar, A.V., Sussman, E.S., Giardino, W.J., and Halpern, C.H. (2018). The nucleus accumbens and alcoholism: A target for deep brain stimulation. Neurosurg. Focus 45, 1–10. 10.3171/2018.5.FOCUS18157.

35. Miller, K.J., Müller, K.R., and Hermes, D. (2021). Basis profile curve identification to understand electrical stimulation effects in human brain networks. PLOS Comput. Biol. 17, e1008710. 10.1371/JOURNAL.PCBI.1008710.

36. Renier, N., Wu, Z., Simon, D.J., Yang, J., Ariel, P., andTessier-Lavigne, M. (2014). iDISCO: a simple, rapid method to immunolabel large tissue samples for volume imaging. Cell 159, 896–910. 10.1016/J.CELL.2014.10.010.

37. Berg, S., Kutra, D., Kroeger, T., Straehle, C.N., Kausler, B.X., Haubold, C., Schiegg, M., Ales, J., Beier, T., Rudy, M., et al. (2019). Ilastik: Interactive Machine Learning for (Bio)Image Analysis. Nat. Methods 16, 1226–1232. 10.1038/s41592-019-0582-9.

38. Wu, H., Miller, K.J., Blumenfeld, Z., Williams, N.R., Ravikumar, V.K., Lee, K.E., Kakusa, B., Sacchet, M.D., Wintermark, M., Christoffel, D.J., et al. (2018). Closing the loop on impulsivity via nucleus accumbens delta-band activity in mice and man. Proc. Natl. Acad. Sci. U. S. A. 115, 192–197. 10.1073/PNAS.1712214114/SUPPL_FILE/PNAS.1712214114.SM02.AVI.

39. Rijsketic, D.R., Casey, A.B., Barbosa, D.A.N., Zhang, X., Hietamies, T.M., andRamirez-ovalle, G. (2023). UNRAVELing the synergistic effects of psilocybin and environment on brain-wide immediate early gene expression in mice. 10.1038/s41386-023-01613-4.

40. Perens, J., Salinas, C.G., Skytte, J.L., Roostalu, U., Dahl, A.B., Dyrby, T.B., Wichern, F., Barkholt, P., Vrang, N., Jelsing, J., et al. (2021). An Optimized Mouse Brain Atlas for Automated Mapping and Quantification of Neuronal Activity Using iDISCO+ and Light Sheet Fluorescence Microscopy. Neuroinformatics 19, 433–446. 10.1007/s12021-020-09490-8.

41. Goubran, M., Leuze, C., Hsueh, B., Aswendt, M., Ye, L., Tian, Q., Cheng, M.Y., Crow, A., Steinberg, G.K., McNab, J.A., et al. (2019). Multimodal image registration and connectivity analysis for integration of connectomic data from microscopy to MRI. Nat. Commun. 10, 1–17. 10.1038/s41467-019-13374-0.

42. Huang, H., Gregg, N.M., Valencia, G.O., Brinkmann, B.H., Lundstrom, B.N., Worrell, G.A., Miller, K.J., and Hermes, D. (2022). Electrical stimulation of temporal and limbic circuitry produces distinct responses in human ventral temporal cortex. bioRxiV.

43. Mitzdorf, U. (1985). Current source-density method and application in cat cerebral cortex: Investigation of evoked potentials and EEG phenomena. Physiol. Rev. 65, 37–100. 10.1152/physrev.1985.65.1.37.

44. Cho, S., Hachmann, J.T., Balzekas, I., In, M.H., Andres-Beck, L.G., Lee, K.H., Min, H.K., and Jo, H.J. (2019). Resting-state functional connectivity modulates the BOLD activation induced by nucleus accumbens stimulation in the swine brain. Brain Behav. 9, 1–17. 10.1002/brb3.1431.

45. Albaugh, D.L., Salzwedel, A., Van DenBerge, N., Gao, W., Stuber, G.D., and Shih, Y.Y.I. (2016). Functional Magnetic Resonance Imaging of Electrical and Optogenetic Deep Brain Stimulation at the Rat Nucleus Accumbens. Sci. Rep. 6, 1–13. 10.1038/srep31613.

46. Dragunow, M., and Faull, R. (1989). The use of c-fos as a metabolic marker in neuronal pathway tracing. J. Neurosci. Methods 29, 261–265. 10.1016/0165-0270(89)90150-7.

47. Stark, J.A., Davies, K.E., Williams, S.R., and Luckman, S.M. (2006). Functional magnetic resonance imaging and c-Fos mapping in rats following an anorectic dose of m-chlorophenylpiperazine. Neuroimage 31, 1228–1237. 10.1016/j.neuroimage.2006.01.046.

48. McIntyre, C.C., Grill, W.M., Sherman, D.L., and Thakor, N.V. (2004). Cellular effects of deep brain stimulation: model-based analysis of activation and inhibition. J. Neurophysiol. 91, 1457–1469. 10.1152/JN.00989.2003.

49. Xu, W., Russo, G.S., Hashimoto, T., Zhang, J., and Vitek, J.L. (2008). Subthalamic nucleus stimulation modulates thalamic neuronal activity. J. Neurosci. 28, 11916–11924. 10.1523/JNEUROSCI.2027-08.2008.

50. Vitek, J.L. (2002). Mechanisms of deep brain stimulation: Excitation or inhibition. Mov. Disord. 17, S69–S72. 10.1002/MDS.10144.

51. Gehrlach, D.A., Weiand, C., Gaitanos, T.N., Cho, E., Klein, A.S., Hennrich, A.A., Conzelmann, K.K., and Gogolla, N. (2020). A whole-brain connectivity map of mouse insular cortex. Elife 9, 1–78. 10.7554/ELIFE.55585.

52. McCracken, C.B., and Grace, A.A. (2009). Nucleus accumbens deep brain stimulation produces region-specific alterations in local field potential oscillations and evoked responses in vivo. J. Neurosci. 29, 5354–5363. 10.1523/JNEUROSCI.0131-09.2009.

53. Nakae, T., Matsumoto, R., Kunieda, T., Arakawa, Y., Kobayashi, K., Shimotake, A., Yamao, Y., Kikuchi, T., Aso, T., Matsuhashi, M., et al. (2020). Connectivity Gradient in the Human Left Inferior Frontal Gyrus: Intraoperative Cortico-Cortical Evoked Potential Study. Cereb. Cortex 30, 4633–4650. 10.1093/CERCOR/BHAA065.

54. Allen, W.E., DeNardo, L.A., Chen, M.Z., Liu, C.D., Loh, K.M., Fenno, L.E., Ramakrishnan, C., Deisseroth, K., and Luo, L. (2017). Thirst-associated preoptic neurons encode an aversive motivational drive. Science (80-.). 357, 1149–1155. 10.1126/science.aan6747.

55. DeNardo, L.A., Liu, C.D., Allen, W.E., Adams, E.L., Friedmann, D., Fu, L., Guenthner, C.J., Tessier-Lavigne, M., and Luo, L. (2019). Temporal evolution of cortical ensembles promoting remote memory retrieval. Nat. Neurosci. 22, 460–469. 10.1038/S41593-018-0318-7.

56. Smith, S.M., and Nichols, T.E. (2009). Threshold-free cluster enhancement: addressing problems of smoothing, threshold dependence and localisation in cluster inference. Neuroimage 44, 83–98. 10.1016/J.NEUROIMAGE.2008.03.061.

57. Sternberg, S. (1995). Biomedical Image Processing. Australas. Phys. Eng. Sci. Med. 18, 26–38.

58. Schindelin, J., Arganda-Carreras, I., Frise, E., Kaynig, V., Longair, M., Pietzsch, T., Preibisch, S., Rueden, C., Saalfeld, S., Schmid, B., et al. (2012). Fiji: An open-source platform for biological-image analysis. Nat. Methods 9, 676–682. 10.1038/nmeth.2019.

59. Parvizi, J., Veit, M.J., Barbosa, D.A.N., Kucyi, A., Perry, C., Parker, J.J., Shivacharan, R.S., Chen, F., Yih, J., Gross, J.J., et al. (2022). Complex negative emotions induced by electrical stimulation of the human hypothalamus. Brain Stimul. 15, 615–623. 10.1016/j.brs.2022.04.008.

60. Keller, C.J., Honey, C.J., Mégevand, P., Entz, L., Ulbert, I., and Mehta, A.D. (2014). Mapping human brain networks withcortico-ortical evoked potentials. Philos. Trans. R. Soc. B Biol. Sci. 369. 10.1098/rstb.2013.0528.

61. Shine, J.M., and Poldrack, R.A. (2018). Principles of dynamic network reconfiguration across diverse brain states. Neuroimage 180, 396–405. 10.1016/j.neuroimage.2017.08.010.

