## Supplemental Figures for "Cross-species brain-wide mapping reveals a conserved and coordinated network engaged by NAc DBS"

### Supplementary Material

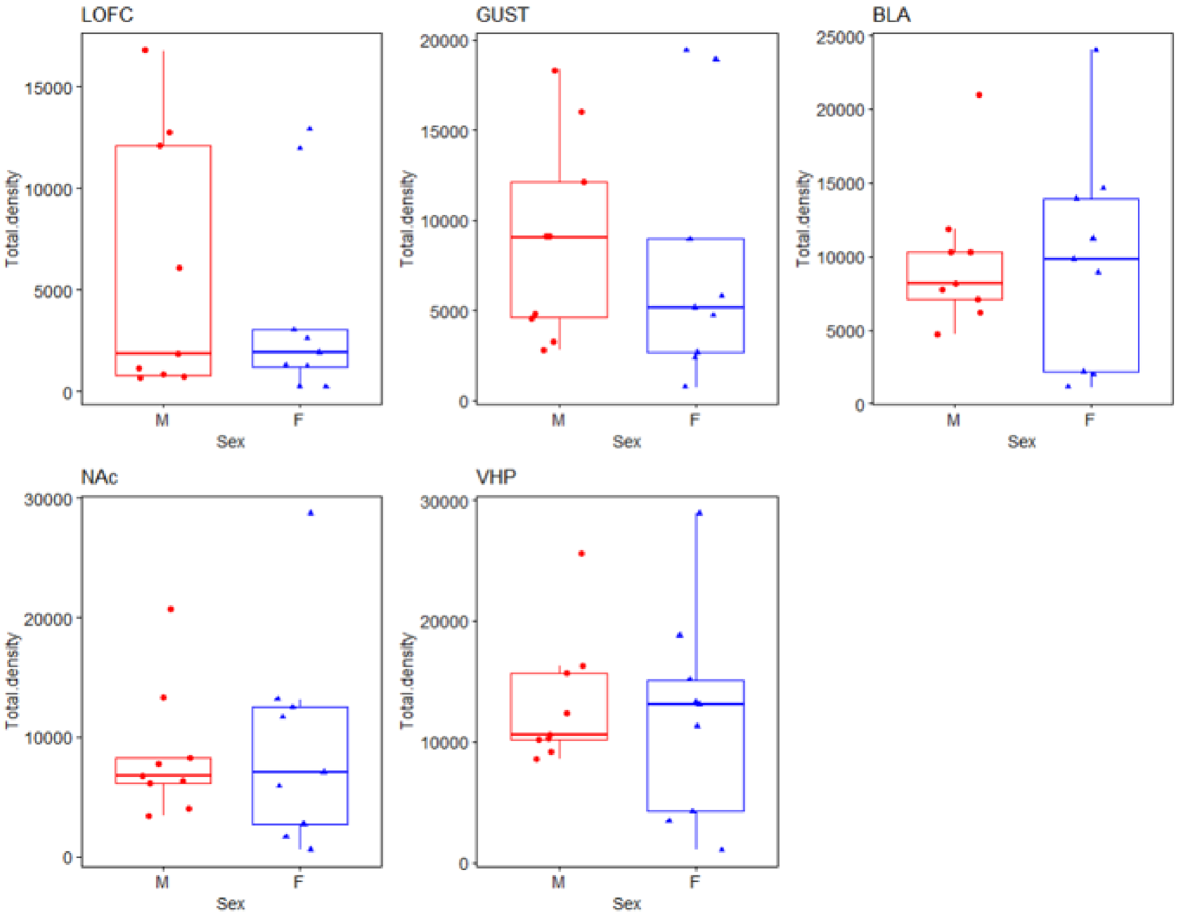


**Supplementary Figure 1. No differences between male female mice stimulation cell counts.**

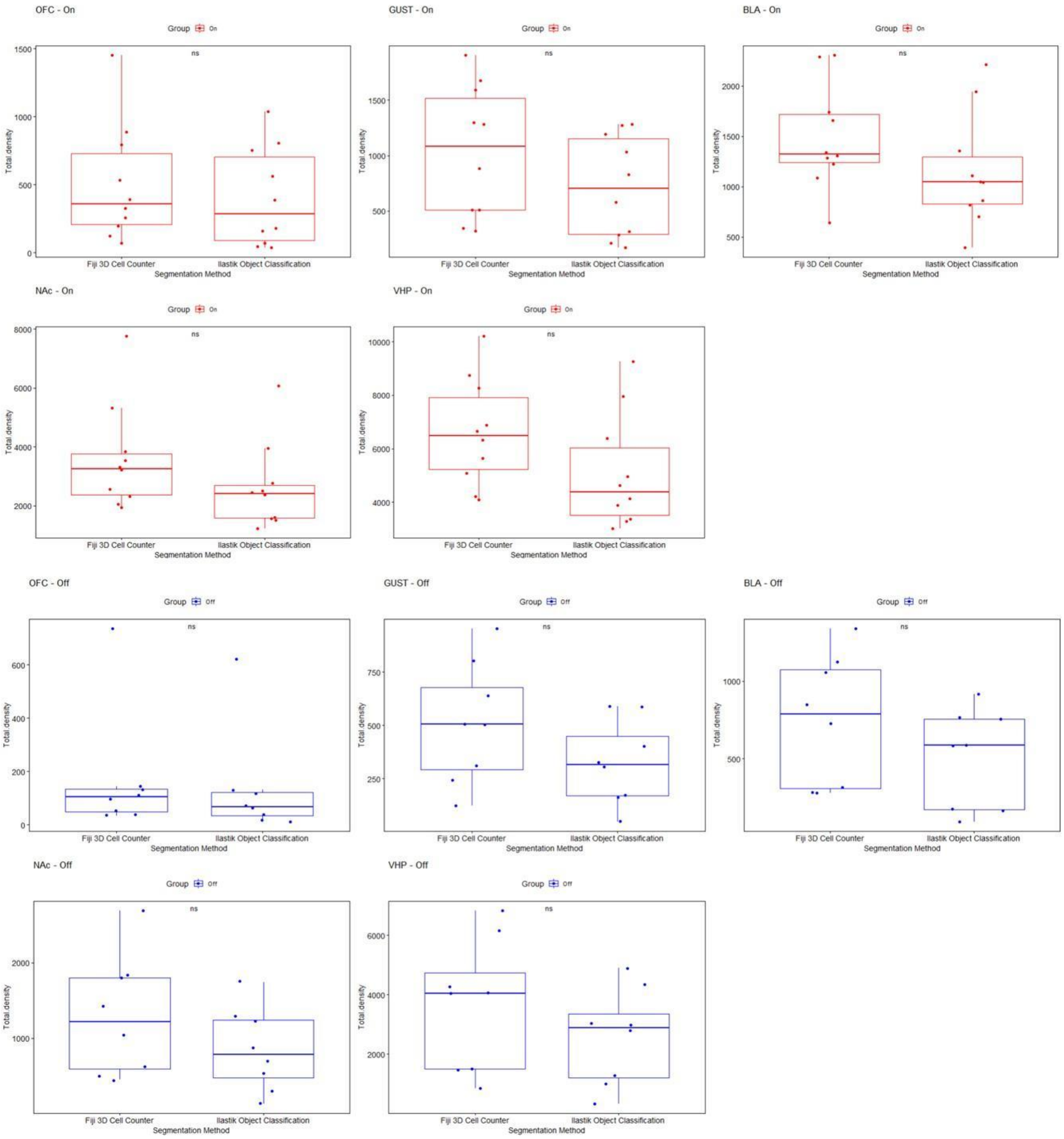


**Supplementary Figure 2. No significant differences between segmentation methods**

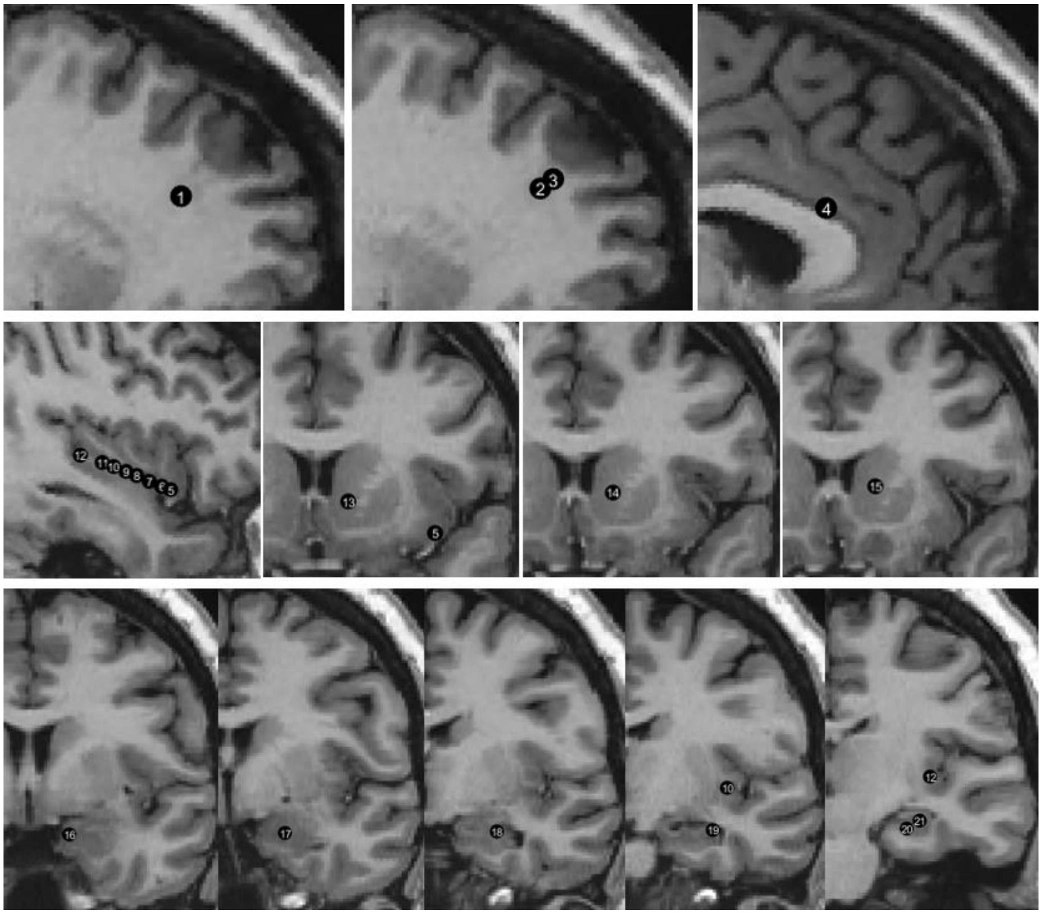


**Supplementary Figure 3. Stimulation sites plotted in corresponding anatomical MRI slices.**

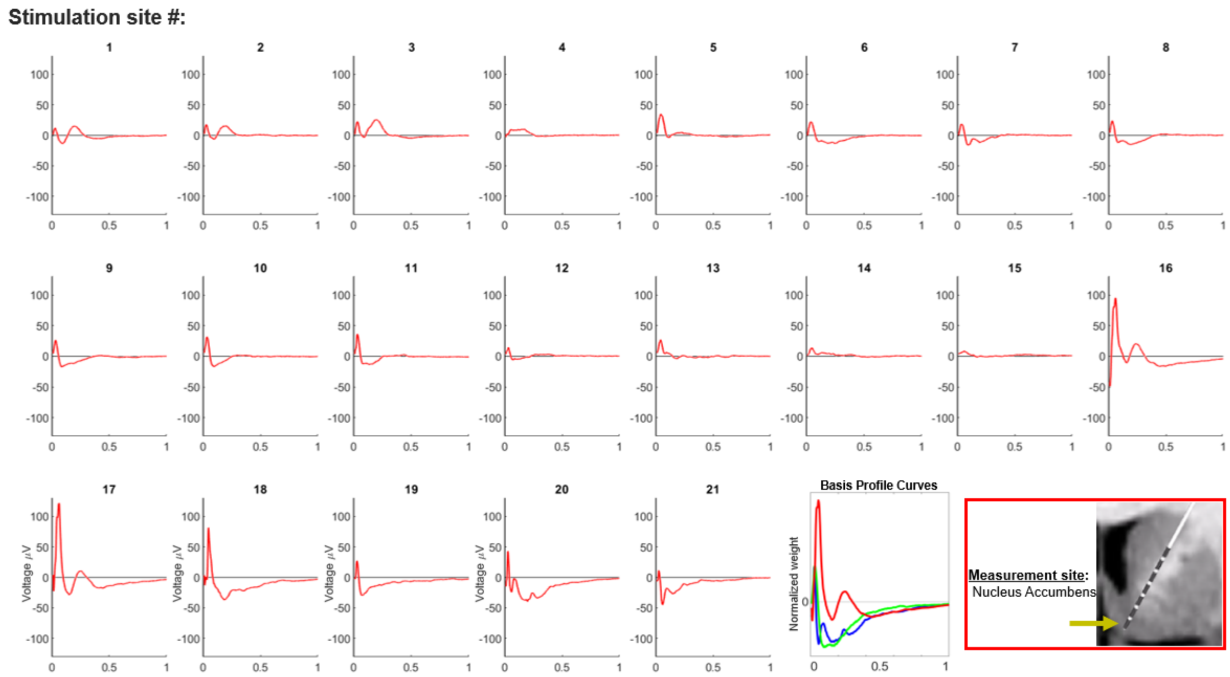


**Supplementary Figure 4. Nucleus Accumbens evoked potentials for each stimulation site.**

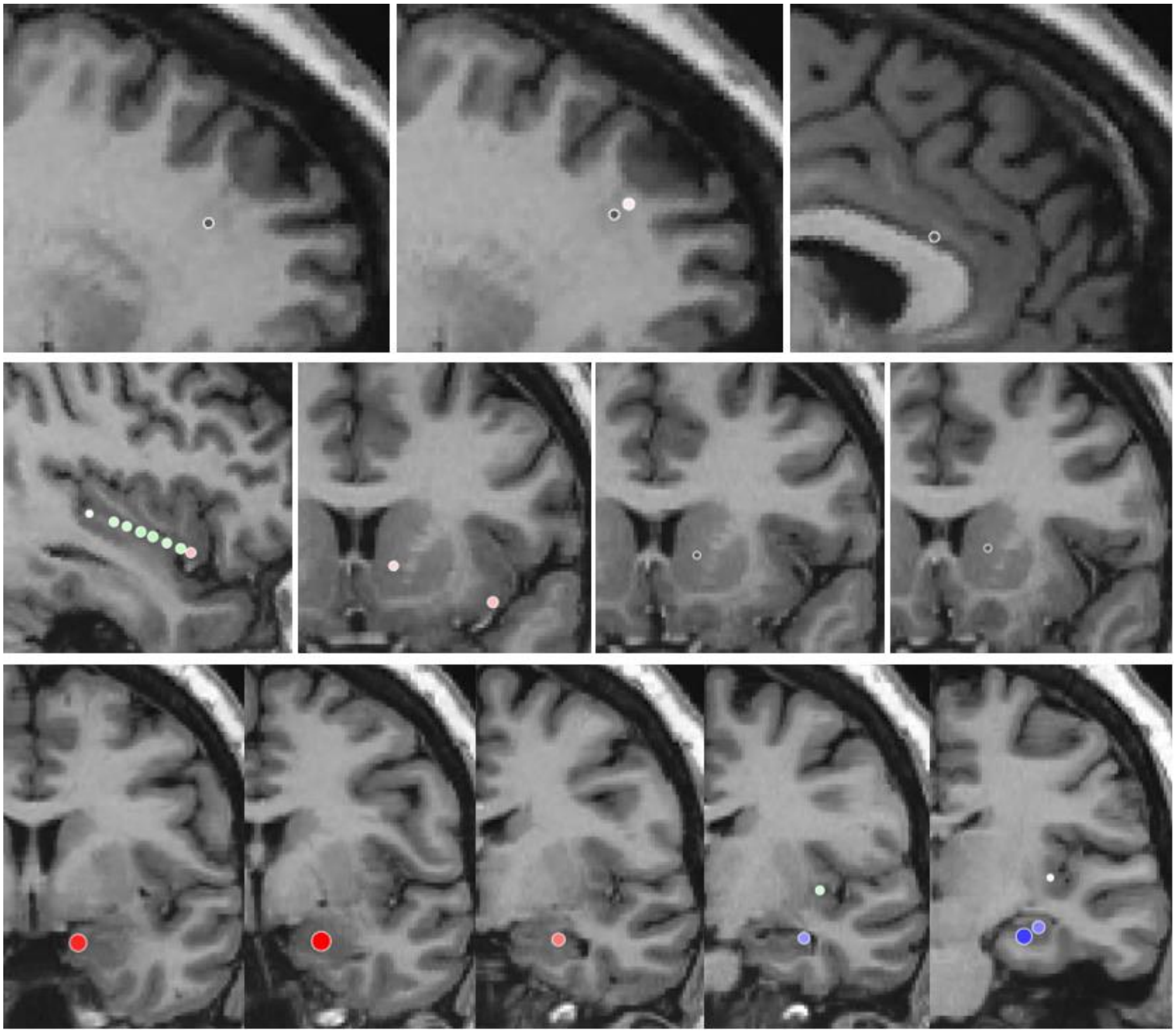

**Supplementary Figure 5. Projection profile and weight plotted in anatomical MRI slices corresponding to stimulation sites.**
